# GenIE-Sys: Genome Integrative Explorer System

**DOI:** 10.1101/808881

**Authors:** Chanaka Mannapperuma, John Waterworth, Nathaniel Street

## Abstract

There are an ever-increasing number of genomes being sequenced, many of which have associated RNA sequencing and other genomics data. The availability of user-friendly web-accessible mining tools ensures that these data repositories provide maximum benefit to the community. However, there are relatively few options available for setting up such standalone frameworks. We developed the Genome Integrative Explorer System (GenIE-Sys) to set up web resources to enable search, visualization and exploration of genomics data typically generated by a genome project.

GenIE-Sys is implemented in PHP, JavaScript and Python and is freely available under the GNU GPL 3 public license. All source code is freely available at the GenIE-Sys website (https://geniesys.org) or GitHub (http://github.com/plantgenie/geniesys.git). Documentation is available at http://geniesys.readthedocs.io.

## Introduction

The availability of tools for setting up standalone genomic web resources is becoming increasingly important given the ever-increasing number of genomes being sequenced. Existing web resources such as Phytozome (1) and Ensembl (2), for example, provide search and navigation of curated genomes but do not allow users to set up their own web resource. While tools for genome browsing exist, these are not designed to serve the data as a public or private web resource. While there are existing options for populating custom resources with genome data, such as the GMOD (Generic Model Organism Database Project) Tripal project (3), BioMart (4) and Intermine (5), we found that these were either more extensive than our own needs or that the available tools did not include visualization of gene expression data. This prompted us to develop an in-house system that was initially used for the PlantGenIE resource(6) and that we have now developed as a generic tool, the Genome Integrative Explorer System (GenIE-Sys) (Supplementary Table 2A). GenIE-Sys is a file-based Content Management System(CMS) (7) with basic content stored in text files. Database server is not needed to get started with the GenIE-Sys. However, the database server is required to load the genomic data and integrate with GenIE-Sys plugins. GenIE-Sys uses the initial source files (General Feature Format (GFF3) and FASTA) generated by genome projects as the core input. These input files are parsed and loaded into a predefined database schema, which is then queried by the web front-end to serve data to the user within a graphical user interface (GUI). The web front-end includes the basic functionalities of a Model Organism Database (MOD); for instance, a genome browser and homology search and visualization. GenIE-Sys provides a dedicated workspace for each species, with a consistent GUI, easy and smooth transition between tools and the ability to save gene lists without login, which represents a lower cognitive load to the user (8). Saved gene lists can be used within the included analysis and visualization tools. We have implemented plugins for visualization and exploration of RNA-Sequencing(9) gene expression datasets, including single gene, gene set or gene co-expression network visualization and functional enrichment analyses. However, a base install, which we refer to as a starter package, includes GeneList, Autocomplete box to speed search of gene IDs, BLAST (10), JBrowse (11) and gene information pages plugins (Supplementary Table 1A).

## Results and Discussion

### System Architecture

The GenIE-Sys can be easily explained using a standard three tier architecture comprising the storage, application and presentation layers. It is a client-server architecture in which the logic for the application, data storage and user interface are developed and maintained as independent modules on separate layers. This enables developers to update the stack of one tier without affecting other areas of the application. Therefore, different development teams can work on different areas of the application scaling up or out the application separately at each tier, aiding ease of maintenance.

The storage layer consists of different types of databases, source files and corresponding indices. In the GenIE-Sys, information is parsed from the database to the application layer for processing, and then eventually to the user via the presentation layer. The application layer is the communication media between the presentation and database layers using web services, which contains two components; one for the inbuilt modules, and other one for the external plugins. Inbuilt modules include the essential functions for providing “on the fly” calculations (server-side algorithms calculate the results depending on the query request without pre-storage in a database) and information processing. The external plugins include all the external tools to integrate with GenIE-Sys, for example JBrowse (11), BLAST (10), exImage (expressionImage), exNet, exHeatmap. The presentation layer is based on web 2.0 standards, including AJAX (Asynchronous JavaScript and XML) (12) and HTML (HyperText Markup Language)(13). The presentation layer directly interacts and communicates with end users. We developed critical functionalities of GenIE-Sys themes using an iterative design process comprising usability evaluation methods, including user-based surveys and observations (14).To negate the requirement of user login we deploy a fingerprint system developed by the Electronic Frontier Foundation that has better accuracy than session IDs or IP addresses (15) (16). However, we will also make use of cookies as an alternative approach to uniquely identifying users as the use of such fingerprint methods is being phased out.

The Number of dependencies, frameworks and libraries were minimized when we initiated the GenIE-Sys to make a base install as simple as possible for the maintainers while providing a usable resource for the end users interacting with the site. This will allow maintainers and developers to create new tools and convert existing tools into plugins easily without conflicting with multiple libraries and frameworks. Figure 1 shows the central system architecture of GenIE-Sys. Necessary gene annotations are stored in primary tables using transcript ID as the primary key while expression, genomic and other analysis data are stored in secondary tables. Users can search for genes of interest or keywords within the GeneList tool and then save the results as a separate, named gene list for use in the analysis and visualization tools. User-specific unique IDs enable users to revisit the site and access their previously created set of gene lists. There are several web services that query the databases in order to extract appropriate data for use in plugins available in a base install, such as GeneList, Autocomplete, Gene information pages, BLAST and JBrowse. Optional plugins such as the expression visualization tools exNet, exImage and exHeatmap, which we have previously described (6), can be downloaded and integrated into a GenIE-Sys installation.

**Figure 1.**
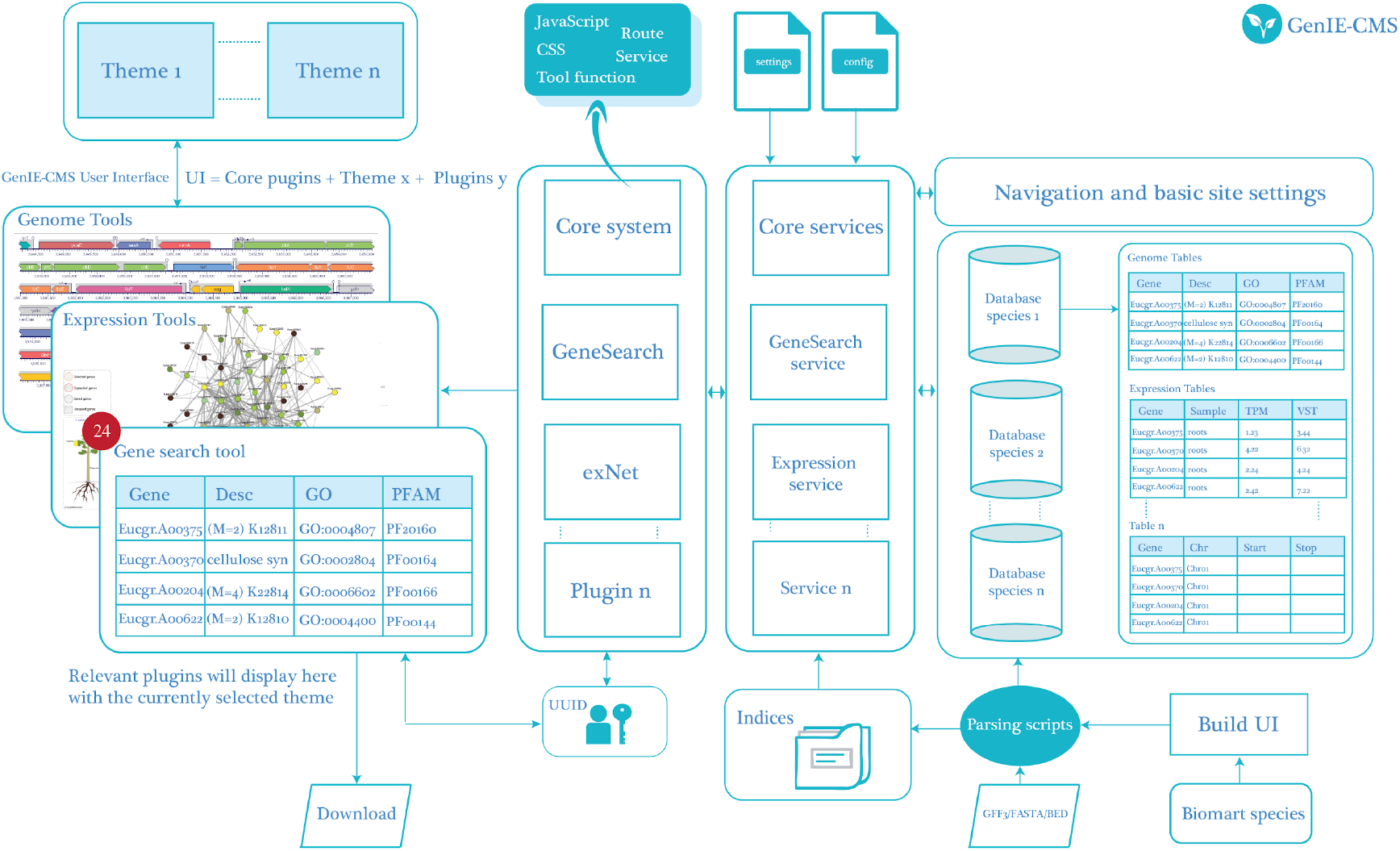
System Architecture of the GenIE-Sys. The core modules are the skeleton of the GenIE-Sys, and it handles the layout properties, directory structure and basic admin features. Default plugin directory contains routing and tool pages plus directories for CSS, JavaScript, and Web Services. Administrators can update the sequence directory paths and MySQL database information inside the config.php file and the settings.php files respectively to configure the GenIE-Sys. Species that are available in Phytozome Biomart can be downloaded using build.plantgenie.org extraction tool to use as input files for parsing scripts. These parsing scripts will be used to load input source files (GFF3/TSV/SQL/FASTA) into the database to make corresponding indices. Separate species stored in a different database.

We have adapted analysis, expression and genome tools from PlantGenIE (6) as external plugins for optional use within the GenIE-Sys.

### Database Architecture

Chado (17) is popular relational database schema covering a variety of biological domains. It is an open-source, community-derived database schema for PostgreSQL. For the time being, there is no associated Chado database for MySQL. Before we introduce the GenIE-Sys, we had to keep two database servers; MySQL for the PlantGenIE(6) tools and PostgreSQL for the Chado database.. Therefore, we had to maintain two separate database servers for a single web resource, and cross-connection between databases is challenging as they are in different database servers. To ease maintenance, we prefer to extract sequence information as BLAST(10) or JBrowse (11) indices instead of parsing and storing them within the database. We already had a well-functioning database structure in PlantGenIE(6) with unique specifications. For example; separate tables for co-expression networks and to be able to save a genelist in the database etc. On the other hand, most of the tables in the Chado database not used in the GenIE-Sys. Therefore, we developed a dedicated MySQL database schema (Supplementary Figure 1A) for GenIE-Sys, which was capable of accomplishing our primary goal to develop tools for gene annotation and expression data visualization. We use transcript gene IDs as primary keys in the database and then combine secondary tables with relation to those primary keys. Therefore, the database structure can be extendable for unforeseen future requirements.

### Navigation system of GenIE-Sys

The data flow diagram (Figure 2) helps to understand the main functionalities and navigation system of the GenIE-Sys.

**Figure 2.**
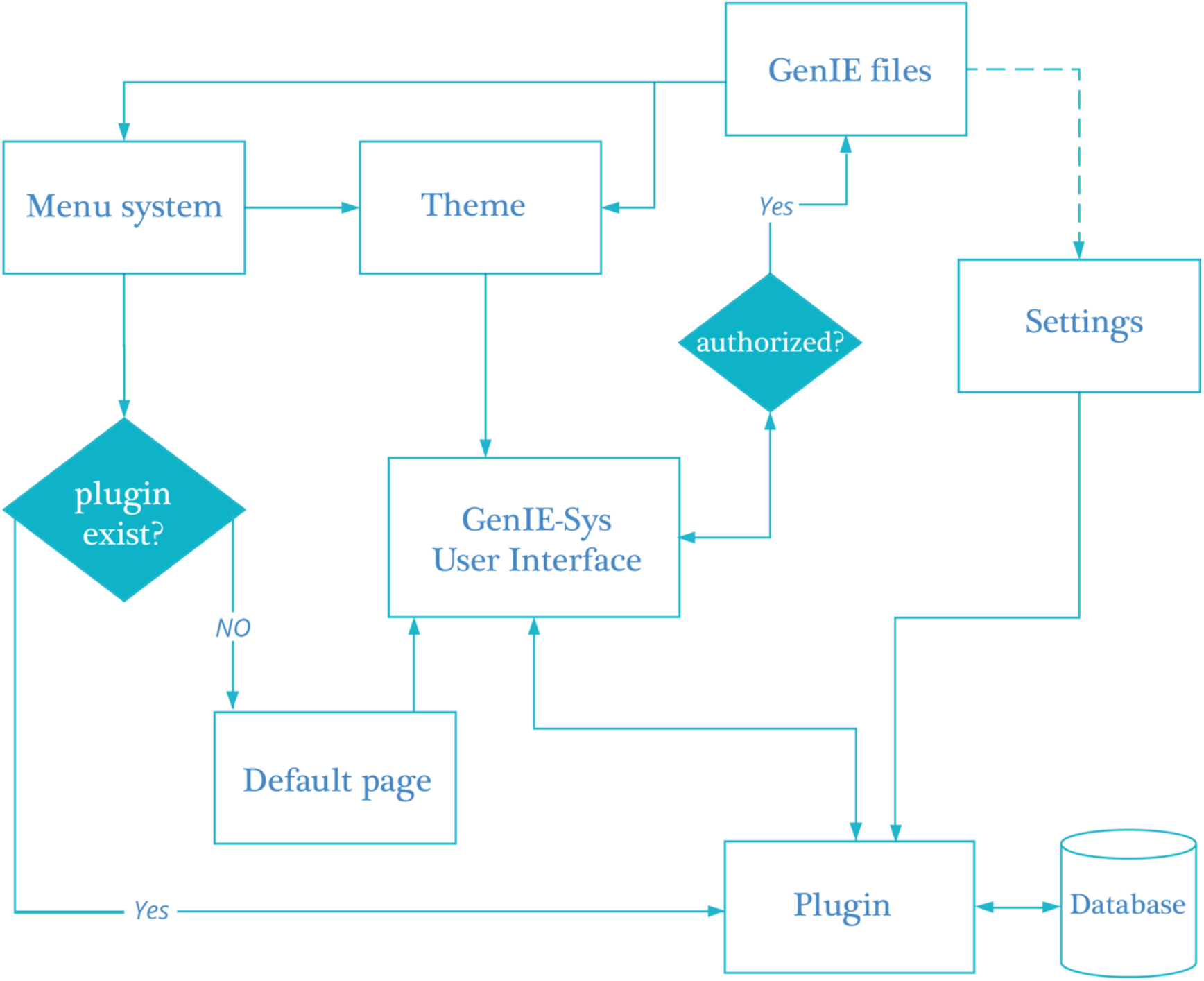
GenIE-Sys Data flow diagram. Genie files can be changed by the administrator using GenIE-Sys user interface. Basic site configurations including passwords, menus and themes information stored in the GenIE files. First, check the availability of the plugin for the given menu name. If the plugin is available for the given menu name, the equivalent plugin will be displayed; otherwise, the user will only see the default blank page. Plugins will use the Database and storage files to visualize genomic, analysis and expression data. The corresponding plugins will render inside the GenIE user interface with the combination of the selected theme. The plugin will also have access to the settings files to get database name, username, password, blast directories and other similar settings. Settings have to access the configuration files to get default database name unless if it’s not already mentioned in the settings files.

### Administration

GenIE-Sys is designed with ease of use and only simple hardware and software requirements in mind. GenIE-Sys can be installed in different infrastructures such as XAMP/MAMP, Docker and Command-line environments. At first, Clone the GenIE-Sys source code from GitHub (https://github.com/plantgenie/geniesys.git) into the Webroot of your web server and website can be access using a web browser. GenIE-Sys is a file-based CMS with essential content stored in text files. MySQL database is required only to load the genomic data and integrate with GenIE-Sys plugins. Once GenIE-Sys is up and running, the administrator can download and install an empty database or model species database using a command-line or graphical user interfaces. Installation requires modest computer skills. Bash scripts are available for parsing genomes into required input formats with the ability to extract data from the Biomart resource (REF) and also for loading data into the databases. Directory of tools is considered as plugins. A plugin is a simple folder containing a collection of PHP files. index.php file is a template which validates the necessary conditions whether the tool is enabled or requested by the menu system and availability of the requested tool. Once conditions are met, tool.php file will be redirected by the index.php file to render the appropriate tool within the web resource. Tool.php contains all the function associated with a particular tool, but the index.php file remains unchanged in any given plugin. Documentation (http://geniesys.readthedocs.io/) detailing plugin development and guidelines are available to build and maintain the GenIE-Sys and develop external plugins. Themes help to style the web resource, and there is a separate theme directory in GenIE-Sys. It can be customized depending on the end-user requirements. To change or update the themes, the administrator needs to edit the corresponding style sheet or theme file. There are two basic configuration files one for the primary site setting: the database details and the other for sequence information and raw file locations.

### End User Interaction

To provide a worked example of using the GenIE-Sys we here (Figure 3) use the beta version of our conifer genome resource, ConGenIE (https://beta.congenie.org). A user can start by performing a Gene Search to search for genes using gene IDs, descriptions, experiments, GO IDs and different annotations and then save the result as a new GeneList that can be named and subsequently used with available analysis and visualization tools.

**Figure 3.**
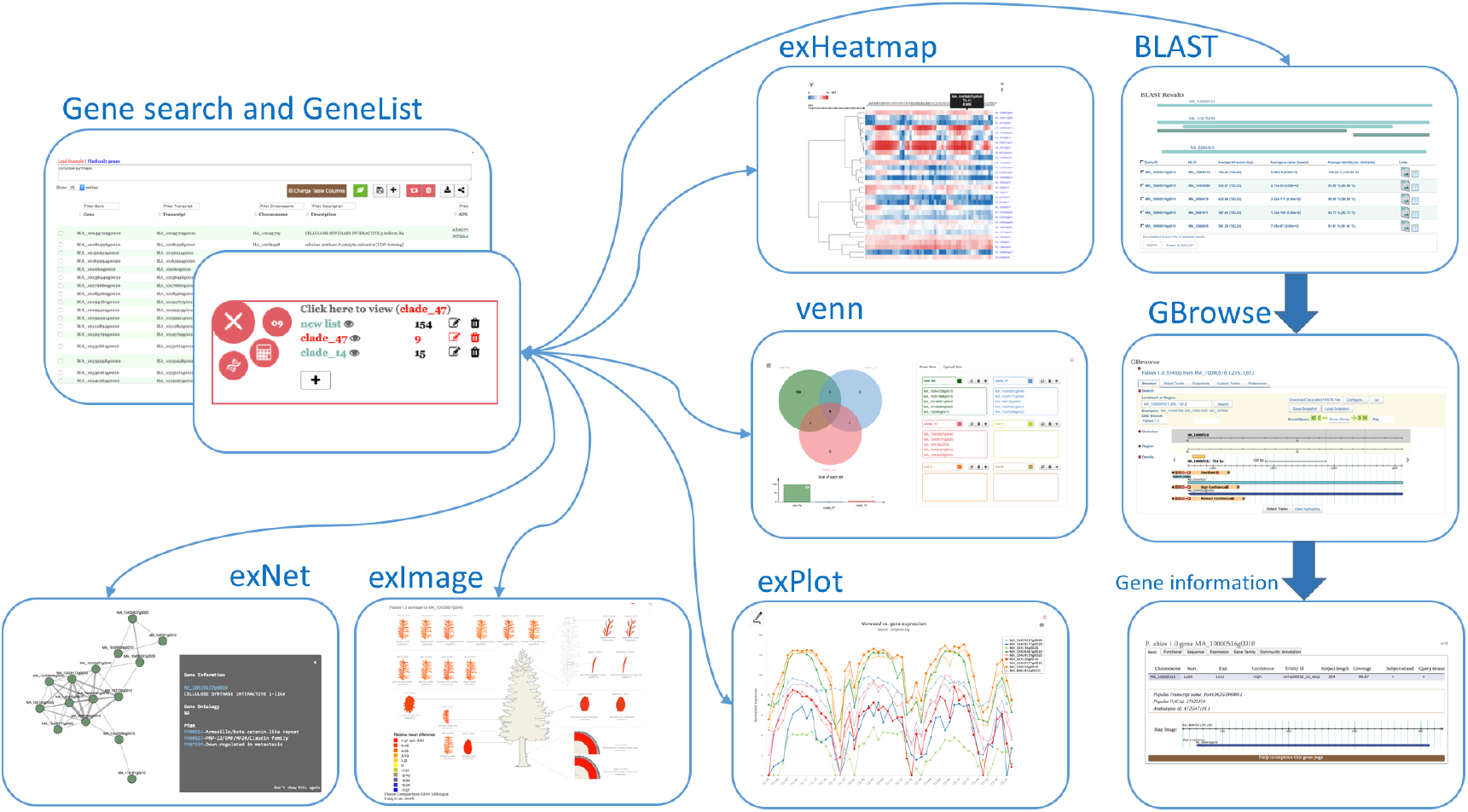
An example of a use case scenario demonstrating beta version of Spruce website (beta.congenie.org). The GeneList tool allows the user to search for genes using gene IDs, descriptions, experiments, GO ids and different annotations, and then saves the result in a list that can be used by other tools. User can type in gene ids, descriptions, or different annotations. The matching genes will be displayed with selected annotations. The result can be customized by clicking the Select Displayed Annotations button. There are three buttons to “Save all to Gene List”, “Remove selected from Gene List” or “Empty Gene List”. The “Share table” button allows sharing the current GeneList with other users by way of an auto generated URL. The GeneList tool is the starting point for most PlantGenIE workflow. User can use saved genelist with PlantGenIE (6) tools such as exNet, exImage, exHeatmap and explot are considered as expression tools.

### FAIR principles for scientific data management

The vast amounts of scientific data being produced today can no longer be effectively interpreted, communicated and shared through papers. Without proper stewardship of data, potential scientific progress is more and more impeded by fragmented patches of information that could not talk to each other. To harness the power of big and complex data people and machines need to work together. To make this happen the revolution in data production needs of revolution in data stewardship. Good data stewardship follows FAIR guidelines(18) and will enable an Internet of data and services helping data and algorithms to find each other talk to each other and remain available for re-use. Data stewardship is, therefore, critical for the future of research and innovation. GenIE-Sys contribute to such efforts as detailed in Table 1.

**Table 1.** GenIE-Sys implemented based on FAIR guidelines

| **FAIR principles** | **GenIE-Sys features** |
| --- | --- |
| **Findable** | Available on GitHub with free and open source License. GenIE-Sys uses consistent layout and workflow for genomic web resources. It also uses standard representation such as gene ids, synonyms for searching. |
| **Accessible** | Once setup the GenIE-Sys anyone can access as a resource unless if data is not private. It provides long term availability. |
| **Interoperable** | GenIE-Sys uses standard files as input source such as GFF3 or FASTA files from the projects. Cross site communication can be achieved when there are more than one species installed. |
| **Reusable** | Results of the tools is reproduceable using corresponding gene IDs. Moreover, shared gene list can be reused among researchers or publications. |

## Conclusion

GenIE-Sys is a standalone web resource builder to produce and integrate a set of tools to explore and interact with genomics data allowing maximal data access and sharing among project members, including those with limited programming skills. Alternatively, the website generated from GenIE-Sys can be publicly accessible as a community resource, ensuring that genome project data is available for broader community use.

## Future

We are planning to continuously update and maintain the GenIE-Sys by developing more features and plugins as the need for these from users is identified. Frequent testing will be performed in order to assure and maintain quality and to deliver bug-free and secure software to the community. Meanwhile, the source code will be revised and re-implemented using new frameworks such as React (20) or Angular (21) to aid community contribution and interpretability. We also aim to developed new tools and plugins for specific biological datasets such as expression and phenotype quantitative trait loci (QTL) and genome wide association studies (GWAS), single cell gene expression data, multi-omics studies and metatranscriptomics and genomics results. Data preprocessing and administration panel along with an API will be further improved into the GenIE-Sys as a fully functional Content Management System to allow researchers to easily upload, modify and maintain data. Currently, there is no security testing or group level authentication; this can be another update by adding a layer to the GenIE-Sys. We believe the development of GenIE-Sys will be a collaborative effort in the future. Hence, we published our source code under the GNU public license on GitHub.

## Authors’ contributions

CM and NR carried out the development, testing and drafted the manuscript. JW helped to conduct usability studies and drafted the final manuscript.

## Acknowledgements

We thank members of the UPSC Bioinformatics platform for informative discussion and testing. We thank Zander Myburg, Nanette Christie and Karen van der Merwe for testing and feedback. We thank the open source community. (Tripal, Galaxy, Taverna and Ergatis) all previous attempt of the genomic databases and online tools. NRS and CM are supported by the Trees for the Future project (T4F).

## References

1. Goodstein DM, Shu S, Howson R, Neupane R, Hayes RD, Fazo J, et al. Phytozome: a comparative platform for green plant genomics. Nucleic Acids Res [Internet]. 2012 Jan;40(Database issue):D1178–86. Available from: http://www.ncbi.nlm.nih.gov/pmc/articles/PMC3245001/

2. Hubbard T, Barker D, Birney E, Cameron G, Chen Y, Clark L, et al. The Ensembl genome database project. Nucleic Acids Res [Internet]. 2002;30(1):38–41. Available from: http://dx.doi.org/10.1093/nar/30.1.38

3. Ficklin SP, Sanderson L-A, Cheng C-H, Staton ME, Lee T, Cho I-H, et al. Tripal: a construction toolkit for online genome databases. Database (Oxford). 2011;2011:bar044.

4. Smedley D, Haider S, Ballester B, Holland R, London D, Thorisson G, et al. BioMart - Biological queries made easy. BMC Genomics. 2009;10:1–12.

5. Smith RN, Aleksic J, Butano D, Carr A, Contrino S, Hu F, et al. InterMine: A flexible data warehouse system for the integration and analysis of heterogeneous biological data. Bioinformatics. 2012;28(23):3163–5.

6. Sundell D, Mannapperuma C, Netotea S, Delhomme N, Lin Y-C, Sjödin A, et al. The Plant Genome Integrative Explorer Resource: PlantGenIE.org. New Phytol [Internet]. 2015 Dec;208(4):1149–56. Available from: http://dx.doi.org/10.1111/nph.13557

7. Rockley A, Kostur P, Manning S. Managing Enterprise Content: A Unified Content Strategy. 2003;

8. Pilar DR, Jaeger A, Gomes CFA, Stein LM. Passwords Usage and Human Memory Limitations: A Survey across Age and Educational Background. PLoS One. 2012;7(12).

9. Mortazavi A, Williams BA, McCue K, Schaeffer L, Wold B. Mapping and quantifying mammalian transcriptomes by RNA-Seq. Nat Methods [Internet]. 2008;5. Available from: http://dx.doi.org/10.1038/nmeth.1226

10. Altschul SF, Gish W, Miller W, Myers EW, Lipman DJ. Basic local alignment search tool. J Mol Biol [Internet]. 1990 Oct;215(3):403–10. Available from: http://linkinghub.elsevier.com/retrieve/pii/S0022283605803602

11. Skinner ME, Uzilov A V, Stein LD, Mungall CJ, Holmes IH. JBrowse: A next-generation genome browser. Genome Res [Internet]. 2009 Sep;19(9):1630–8. Available from: http://www.ncbi.nlm.nih.gov/pmc/articles/PMC2752129/

12. Garrett JJ. Ajax: A New Approach to Web Applications by. Semin Respir Infect. 2005;1(2):1–5.

13. HTML.

14. Mannapperuma C, Street N, Waterworth J. Designing usable Bioinformatics Tools for specialized users. In: ICITS’19 - The 2019 International Conference on Information Technology & Systems. Quito, Ecuador; 2019.

15. Eckersley P. How Unique is Your Web Browser? In: Proceedings of the 10th International Conference on Privacy Enhancing Technologies [Internet]. Berlin, Heidelberg: Springer-Verlag; 2010. p. 1–18. (PETS’10). Available from: http://dl.acm.org/citation.cfm?id=1881151.1881152

16. Boda K, Földes ÁM, Gulyás GG, Imre S. User Tracking on the Web via Cross-Browser Fingerprinting. In: Laud P, editor. Information Security Technology for Applications. Berlin, Heidelberg: Springer Berlin Heidelberg; 2012. p. 31–46.

17. Mungall CJ, Emmert DB, Gelbart WM, de Grey A, Letovsky S, Lewis SE, et al. A Chado case study: An ontology-based modular schema for representing genome-associated biological information. Bioinformatics. 2007;23(13):337–46.

18. Wilkinson MD, Dumontier M, Aalbersberg IjJ, Appleton G, Axton M, Baak A, et al. Comment: The FAIR Guiding Principles for scientific data management and stewardship. Sci Data. 2016;3:1–9.

19. Manley G. Public Access NIH Public Access. 2013;71(2):233–6.

20. ReactJS.

21. AngularJS.

22. Atanasovska B, Kumar V, Fu J, Wijmenga C, Hofker MH. Review GWAS as a Driver of Gene Discovery in Cardiometabolic Diseases. Trends Endocrinol Metab. 2015;26(12):722–32.

