## Supplementary material for "GenIE-Sys: Genome Integrative Explorer System"

### GenIE-Sys Database Diagram

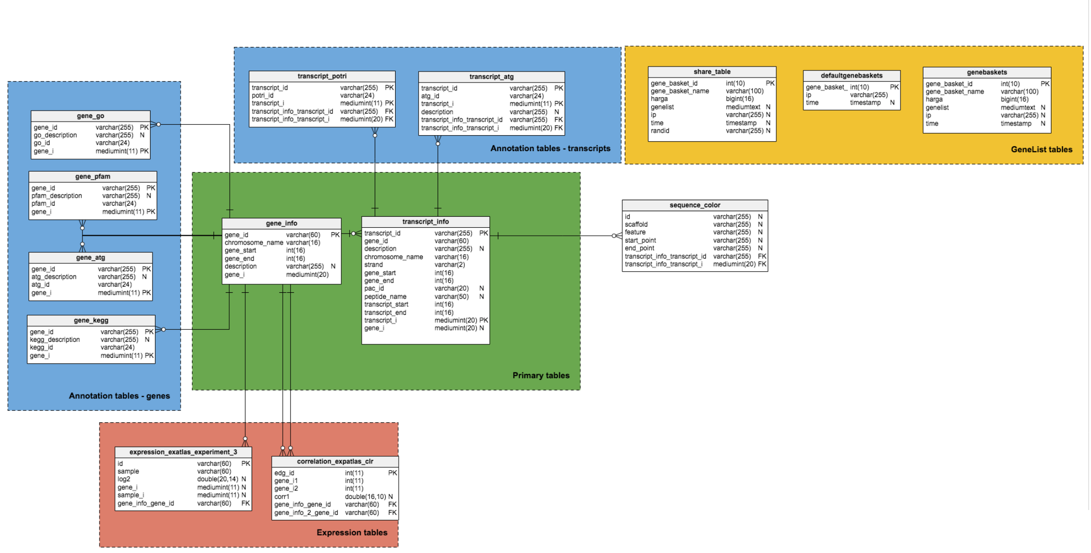


**Figure 1A  Database diagram of GenIE-Sys**

transcript_info and gene_info tables are considered as primary tables and any of the secondary tables can be changed updated or added without affecting the ain system.

**Table 1A Available Plugins in GenIE-Sys**

| Plugin name | Included in Starter package | Dependencies | Source | Data storage |
|---|---|---|---|---|
| GeneList | Yes | Basic | PlantGenIE | Primary |
| Autocomplete | Yes | Basic | PlantGenIE | Primary |
| gene information | Yes | Basic | PlantGenIE | Primary and Secondary |
| BLAST | Yes | NCBI Blastall | NCBI | Primary/ BLAST indices |
| JBrowse | Yes | JBrowse | GMOD | JBrowse indices |

| | | | | |
|---|---|---|---|---|
| exImage | No | Basic | PlantGenIE | Secondary/ expression |
| exNet | No | Cytoscape | PlantGenIE | secondary/ network |
| exHeatmap | No | Basic | PlantGenIE | Secondary/ expression |
| exPlot | No | Basic | PlantGenIE | Secondary/ expression |
| Venn | No | Basic | PlantGenIE | Primary |

*Note*: All the plugins were developed using JQuery, d3js libraries and PHP, JavaScript, HTML programming languages as Basic dependencies.

**Table 2A Comparision with other resources**

| System | Advantages | Disadvantages |
|---|---|---|
| Tripal | -Supports programmatic access via RESTful API<br>-Continuous maintenance and support<br>- User friendly Interface<br>-Multiple user level and permissions | -Tripal heavily depend on Drupal Content Management system. Which means Tripal has to face all the difficulties that Drupal faces (version wars, memory hog, abandoned modules, steep learning curve, e.t.c) as discussed in Supplementary Table 3A.<br>- Steep learning curve to make new tools<br>-GeneLIst can not be saved<br>- Need to have Unix skills to install and maintain the system |
| InterMine | -Parsers for integrating data from nu-merous formats<br>-Web access to integrated data at a number of levels, from simple brows-<br>ing to complex queries<br>-Facilities for adding one's own data<br>-User friendly web interface that can be<br>easily customised<br>-Data provenance is tracked<br>-GeneList can be saved once user login to the system | -No tools for classifying or clustering data<br>-Queries don't include similarity functions to address annotation errors<br>-Need comparatively high level of infrastructure<br>- Need to have Unix skills to install and maintain the system |
| GenIE-Sys | -User Friendly interface to query the database.<br>- Can set up with modest computer | -Unable to create multiple users or user level permission system<br>-Limited number of modules are |

| | | |
|---|---|---|
| | skills<br>-Flexible and extensible database infrastructures<br>-Minimum number of dependencies dependencies<br>-Easy integration of new data and tools<br>-Easily customisable themes, modules and database<br>-Tools build using other frameworks can be easily integrated into GenIE-Sys plugin system.<br>-GeneList can be saved to do further analysis | available<br>-Early stage of the development<br>- Need to have Unix skills to install and maintain the system |
| BioMart | -Tools for federating a variety of bio-logical databases<br>-Unified web-based user-friendly inter-<br>face for data mining<br>-Supports programmatic access (Perl API, RESTful web services)<br>-Queries defined as a set of successive<br>filters | -Queries limited to only two datasets at<br>once<br>-Not possible to edit or create new filters<br>- No option to save genelist<br>- Need to have Unix skills to install and maintain the system |

**Table 3A Implementation of GenIE-Sys**

| **Website** | **Species** | **URL** |
|---|---|---|
| EucGenIE | *Eucalyptusgrandis* v2.0 | https://eucgenie.org/ |
| Beta PopGenIE | *Populus trichocarpa* v3.1,<br>*Populus tremula* v1.1,<br>*Populus tremuloides* v1.0,<br>*Populus tremula X Populus tremuloides* v1.0 | http://beta.popgenie.org |
| Beta ConGenIE | *Picea abies* v1.0<br>*Picea glauca* WS7711<br>*Picea glauca* PG29<br>*Pinus taeda* v1.1 | http://beta.congenie.org |
| Beta AtGenIE | *Arabidopsis thaliana* Araprot 11<br>*Arabidopsis thaliana* TAIR 10 | http://beta.atgenie.org |
| Beta PlantGenIE | *Populus trichocarpa* v3.1,<br>*Populus tremula* v1.1,<br>*Populus tremuloides* v1.0,<br>*Populus tremula X Populus tremuloides* v1.0<br>*Picea abies* v1.0<br>*Picea glauca* WS7711<br>*Picea glauca* PG29<br>*Pinus taeda* v1.1<br>*Arabidopsis thaliana* Araprot 11 | http://beta.plantgenie.org |

| | | |
|---|---|---|
| | *Arabidopsis thaliana* TAIR 10 | |
| YellowHorn | *Xanthoceras sorbifolium* v2.0 | http://yellowhorn.plantgenie.org |
| Zostera marina | *Zostera marina* v2.2 | http://zmarina.plantgenie.org/ |
| Kalyptos at SweTree Technologies | *Eucalyptusgrandis* v2.0 | Private URL |
